# The physiology of bud dormancy and cell cycle status in grapevine

**DOI:** 10.1101/2024.02.22.581692

**Authors:** Dina Hermawaty, Peta L. Clode, John A. Considine, Santiago Signorelli, Michael J. Considine

## Abstract

Evaluating the cell cycle status during dormancy of multicellular organisms is problematic. This is particularly so for woody perennial buds, where dormant and quiescent states are diffuse, and the organ may remain visibly unchanged for six to nine months of the year. In this study, we investigate cell cycle status of dormant grapevine buds by measuring mitotic index using an optimised method developed for grapevine bud tissue. The experimental material showed a dynamic range in the depth of dormancy, declining from 200 days in March to less than 60 days in May and 30 days in August, measured as the time to reach 50% bud burst in forcing conditions. Despite these differences, flow cytometry analysis showed that most nuclei isolated from these buds were arrested at the G1 phase. Ultrastructure analysis of the cells in the region of the shoot apical meristem confirmed that the mitotic activities of buds remained low at all time points, together with the development of starch grains and the relative absence of organelle development.

**HIGHLIGHT:** The cell cycle and ultrastructure data suggest interesting evidence correspond to the growth resumption capacity of grapevine cv. Cabernet Sauvignon buds, i.e., absence of mitosis activities regardless of dormancy depth and starch accumulation irrespective of chilling accumulation.

## INTRODUCTION

The release of dormancy and regulation of bud break has wide economic and ecological importance, as well as scientific interest (Saure, 1985; Heide and Prestrud, 2005; Olsen, 2010; Singh et al., 2017). The role of chilling exposure in dormancy release has become a paradigm, and as a result, much of our knowledge of bud dormancy is restricted during the period of natural chilling exposure, from mid-late autumn through winter (Or et al., 2000; Pérez and Lira, 2005; Vergara and Perez, 2010; Niu et al., 2016; Zheng et al., 2018). Nevertheless, studies in grapevine indicate that bud physiology is highly dynamic during late summer (Pouget, 1963; Nigond, 1967; Velappan et al., 2022). Therein, the depth of dormancy was maximal in late summer, declining substantially prior to the middle of autumn. This study poses a challenge to the classic autumn dormancy and dormancy nomenclature introduced by Lang (1987), suggesting that in some grapevine cultivars the requirement for chilling to be facultative, and its effect is quantitative rather than qualitative (Nigond, 1957).

Since dormancy is a property of meristematic organs, which are defined by the capacity for cell division, we may expect a close relationship between the dynamics of bud dormancy and cell cycle regulation. Several studies have documented changes in the expression of grapevine cell cycle genes, including during developmental and dormancy transitions (Díaz-Riquelme et al., 2012; Vergara et al., 2016 and 2017). However, there is a considerable lack of physiological data available on cell division patterns during the annual growth cycle of proleptic buds, particularly grapevine (Velappan et al., 2017). Earlier observations of mitotic activities in seeds, embryos and terminal buds showed that cells are predominantly arrested at the G1 phase of the cell cycle (see Velappan et al., 2017, Table 1 and references therein). This relationship is well-supported in the case of seed, whereby extreme desiccation restricts metabolic activity (Bewley, 1997). In barley seed having 10-12 % water content, 82 % and 17 % embryo cells were observed at G1 and G2 phase respectively (or at 2C and 4C DNA level), and the G2 proportion rapidly increased to 32 % within 18 hours of imbibition (Gendreau et al., 2012). A recent study observing the water influx into the terminal buds of Norway spruce showed that upon exposure to short-day, a thick-walled cell was formed at the base of the buds preventing water transport into the buds, which coincided with growth cessation and dormant bud formation (Lee et al., 2017). In contrast, the base of quiescent grapevine buds did not appear to be a barrier for water uptake, although a thorough examination throughout dormancy is yet to be made (Signorelli et al., 2020).

Shoot mass emerged from axillary buds during bud burst is generated from cell cycle activity of the meristem at the tip of the shoot primordia, called shoot apical meristem (SAM) (Sussex, 1989). Cells located at the central zone of SAM maintain its pluripotent nature and provide a continuous supply of cells that will differentiate into cells at various organs (Lenhard and Laux, 1999). In most dicot plants, the SAM consists of three layers of cells, namely L1, L2, and L3. In relation to cellular differentiation, the epidermis was found to be exclusively generated from the L1, whereas the inner part of the plant was derived from L2 and L3 cell layers, indicating that cell differentiation in the plant is position dependent not linage dependent (Sussex, 1989; Kaplan and Cooke, 1997; Lenhard and Laux, 1999). Cells at the CZ and its derivatives are mitotically less active than those in peripheral zone (PZ), which contains cells that start the differentiation process (Laufs et al., 1998). In addition to differences in mitotic activities and pluripotent identities of the cells at the SAM, the ultrastructure of cellular components was also found to generate distinct cytological zonation at the shoot primordial (Mayerowitz, 1997; Bowman and Eshed, 2000). At the SAM, nuclear occupy a significant part of the cells as opposed to cells at PZ or rib zone (RZ), which contain large vacuole. Thus, under the microscope, the cells at SAM will appear more dense compared to PZ or RZ. Morphological differences in the organelles are also evident, being organelles at CZ less mature compared to organelles found in the more active zone such as PZ or leaf primordia (LP). Consequently, starch grains and lipid bodies are commonly found in the CZ cells function as an energy reserve and the development of mitochondria are pronounced in the mitotically active cells (Lyndon and Robertson, 1976 and Sawhney et a., 1981). In axillary bud development of pea, the number of plastids remained constant until leaf development was initiated (Lyndon and Robertson, 1976).

Microscopic analysis on terminal buds of *Pseudotsuga menziesii*(Douglas fir) showed that the shoot meristem comprised heterogeneous cells, which differed in size and nuclear DNA content and thus created a histological zonation when observed under the microscope (Owens and Molder, 1972). At the whole shoot apex level, no mitosis was detected during dormancy, with an average of 45 % cells were found to have 2C nuclear DNA content (G1/S in cell cycle phase), and 15 % cells had 4C nuclear DNA content (G2/M). Interestingly, this proportion did not change much, even after the late leaf initiation stage. Furthermore, histological zonation was maintained through bud development and independent of nuclear DNA content. Nevertheless, significant changes can be observed when comparing cell cycle activities at a particular zone along the developmental stage. For instance, the mitotic activities in early bud-scale initiation differ significantly compared to late leaf initiation at the apical zone but not at the peripheral zone, suggesting spatial regulation of mitotic activities.

In this study, we have evaluated cell division status in the primary bud complex of grapevine, comparing stages of dormancy over two consecutive seasons. A quantitative nuclei content measurement using flow cytometry (FCM) and the cellular ultrastructure morphology observation was performed at the shoot apices in intact buds differing in the depth of dormancy. The result indicated that the phenotypic appearance of the bud during dormant stage reflects the cell cycle status and the cellular developmental progression at the SAM.

## RESULTS

### Depth of dormancy

Variation in the time required to achieve 50% budburst among grapevine cultivars is evident in the literature. Pronounced dynamics were reported in cv. Merlot grown in Bordeaux, France (44°N) required more than 100 days to achieve 50% budburst when buds were considered in peak dormancy (Pouget, 1963). Meanwhile, the same cultivar grown in the Margaret River area, Western Australia (34°S) required more than 200 days to attain 50% budburst at peak dormancy stage (Velappan et al., 2022). In this experiment, using cv. Cabernet Sauvignon, measurement to 50% budburst (BB_50_) was conducted up to 300 days. **Figure 1** shows the depth of dormancy of buds collected at the three time points in 2017 (March 5^th^, May 16^th^, August 8^th^) and 2018 (March 5^th^, May 13^th^, August 4^th^). Peak dormancy was observed in the March samples when water-treated buds only achieved 24% bud burst within the 300 days observation period in 2017. The remaining 76% of the water-treated March buds were necrotic. Although our data could not determine whether these buds were necrotic at the time of sampling or became necrotic over the forcing period, the high degree of viability of other samples collected at the same vineyard, including the H_2_CN_2_-treated buds of the same sample date, suggests the latter (**Table 1**). The H_2_CN_2_-treated buds from March achieved BB_50_ within 49 days, and 90% of buds burst within 300 days. In the May samples, water-treated buds reached BB_50_ within 46 days and H_2_CN_2_-treated buds within 18 days. The shortest duration to achieve BB_50_ was observed in the water- and H_2_CN_2_-treated buds collected in August, with 21 and 24 days respectively. The dormancy analysis in 2018, was consistent with 2017 data (**Figure 1** and **Table 1**).

**Figure 1.**
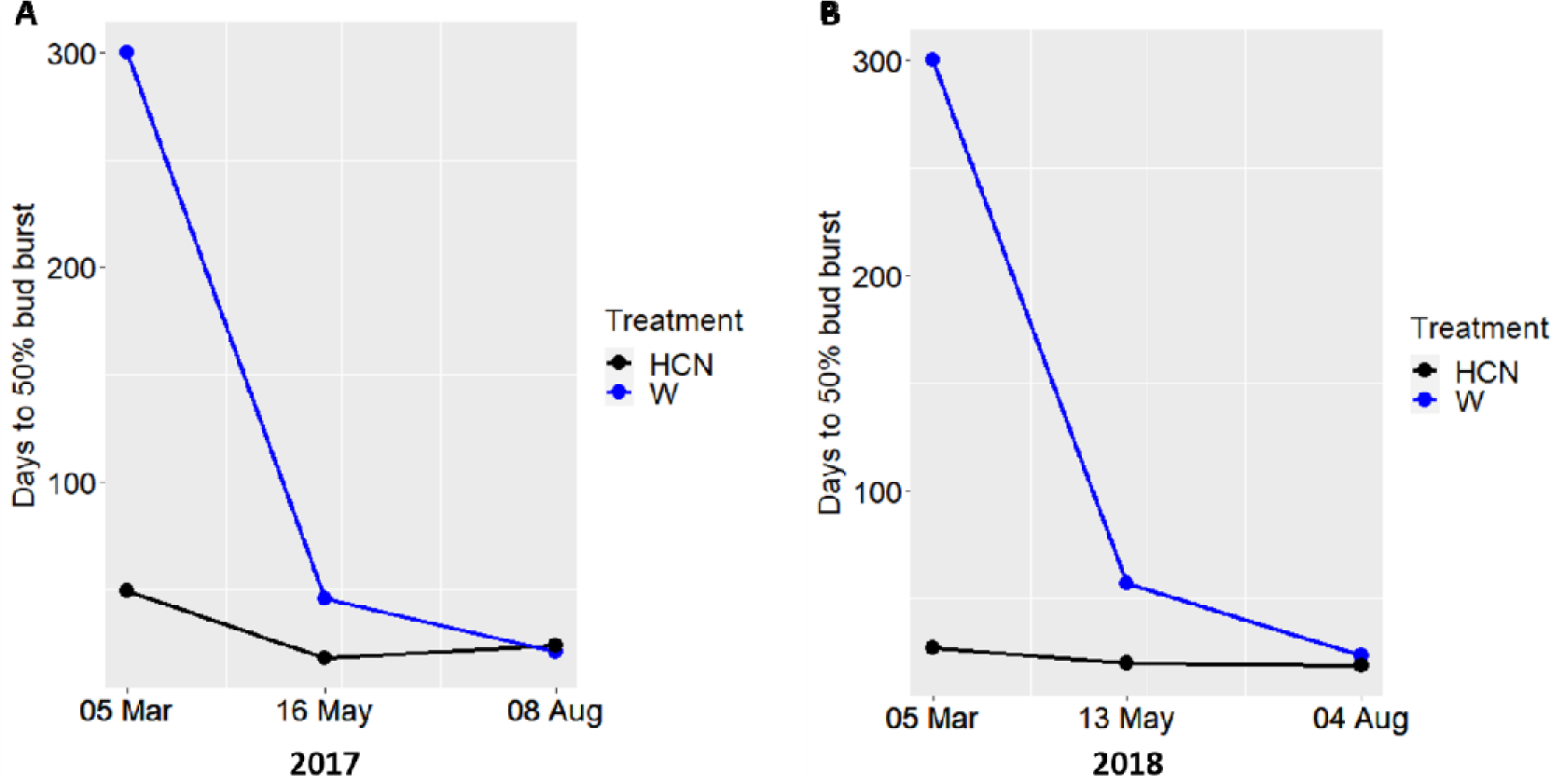
The time to 50% bud burst (BB_50_) of Vitis vinifera cv. Cabernet Sauvignon collected from the Margaret River region in southwest Western Australia (34 °S, 115 °E) in March, May, and August in 2017 (A) and 2018 (B). Days to 50% bud burst of grapevine buds treated with water (blue) and H_2_CN_2_ 1.25% (w/v; black) is presented.

**Table 1.** The number of buds growing into shoot under force growth condition when collected in March, May, and August in 2017 and 2018 and treated with water (W) or hydrogen cyanamide (H_2_CN_2_).

| Harvest Date | Days to 50% bud burst |  | % Bud burst |  |
| --- | --- | --- | --- | --- |
| | W | $\text{H}_2\text{CN}_2$ | W | $\text{H}_2\text{CN}_2$ |
| March 5, 2017 | 365<sup>*</sup> | 49 | 24 | 90 |
| May 16, 2017 | 46 | 18 | 84 | 100 |
| August 8, 2017 | 21 | 24 | 96 | 74 |
| March 5, 2018 | 286<sup>**</sup> | 27 | 32 | 86 |
| May 13, 2018 | 57 | 20 | 96 | 96 |
| August 4, 2018 | 24 | 19 | 100 | 100 |

<sup>\*</sup> 24% bud burst after 150 days and the remaining buds found to be necrotic after 365 days.
<sup>\*\*</sup> 32% bud burst after 175 days and the remaining buds found to be necrotic after 286 days.

### Cell cycle profile of dormant buds

The proportion of cells in each cell cycle phase was estimated from the univariate PI_DNA histogram using the Watson pragmatic algorithm (Watson et al., 1987). Results show that the majority of cells were in the G1 phase of the cell cycle in both years, although there were marginal differences in the magnitude between years (**Table 2**). There was no significant effect of H_2_CN_2_ on the mitotic index within the 24 hours of treatment in either 2017 or 2018 (**Figure 2**). The mitotic index declined significantly from March to May and to August in 2017, and while the trend was consistent in 2018, the differences were not significant. The coefficient of variance (CV) value of the cytometric data and debris level in the nuclei suspension ranged from 10 to 14% and 59% to 79%, respectively (**Table 2**).

**Figure 2.**
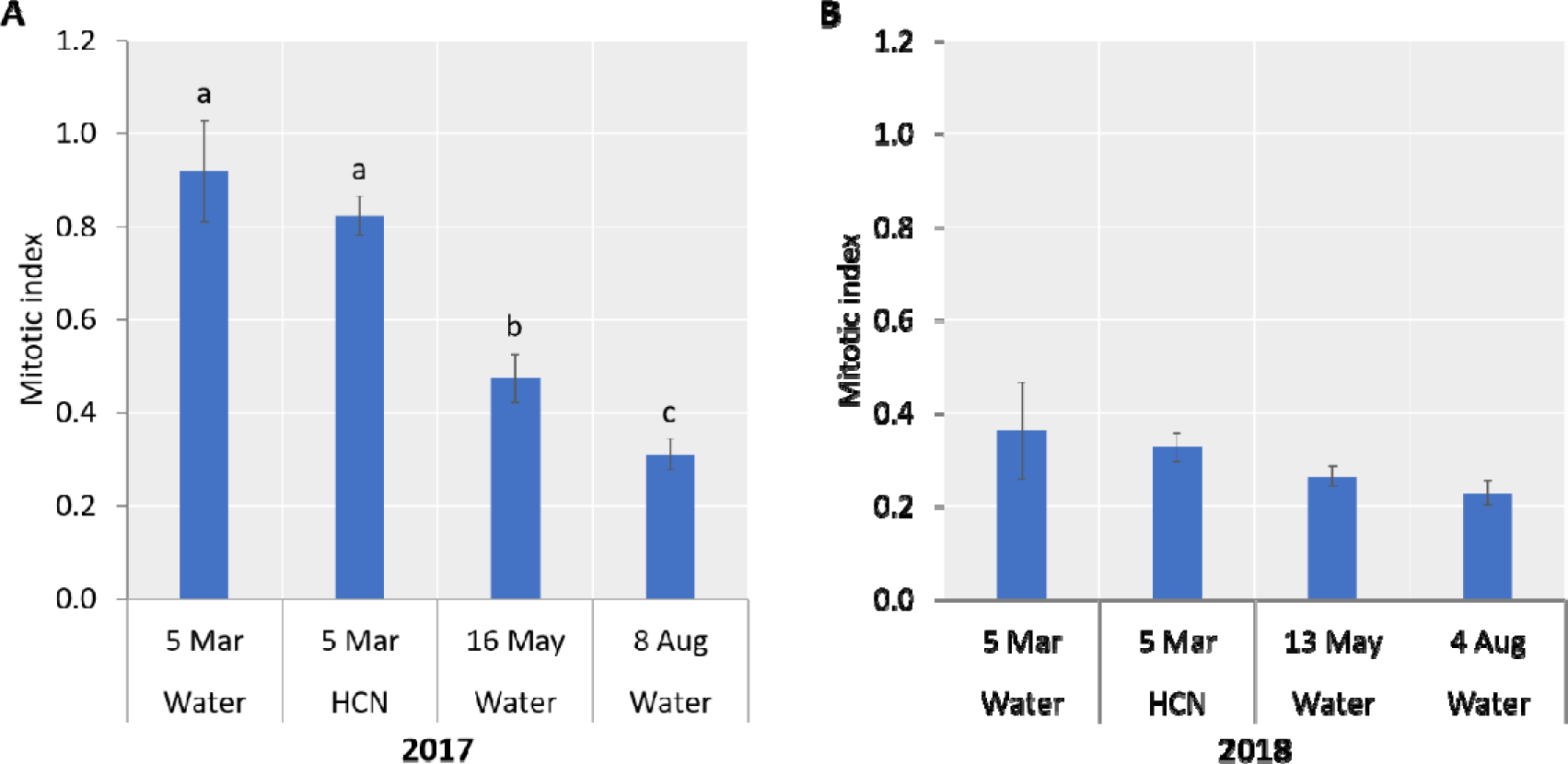
Mitotic index (G2/G1) of nuclei isolated from the primary bud of grapevine buds at each treatment and time in 2017 (**A**) and 2018 (**B**). Different letters indicate significant differences against the respect to the March-Water condition using a Tukey comparison (n = 3, p < 0.05). Trends were consistent between years, but no significant differences were observed between conditions in 2018.

**Table 2.** The proportion of cells in dormant grapevine buds collected in 2017 and 2018 at each cell cycle phase. Treatment was done by submerging buds for 30 seconds in 1.25 % (w/v) hydrogen cyanamide (H_2_CN_2_) solution and water (W).

| Harvest Date | Treatment | G1 | S | G2 | Mitotic index | CV | Debris Factor |
| --- | --- | --- | --- | --- | --- | --- | --- |
| March 5, 2017 | Water | 43.30 ± 6.55 a | 14.31 ± 5.37 ab | 39.33 ± 1.12 c | 0.92 ± 0.11 c | 12.87 ± 1.53 ab | 0.64 ± 0.01 a |
| March 5, 2017 | H<sub>2</sub>CN<sub>2</sub> | 38.63 ± 5.27a | 23.47 ± 7.26 b | 31.80 ± 4.20 b | 0.82 ± 0.04 c | 10.25 ± 2.12 a | 0.59 ± 0.03 a |
| May 16, 2017 | Water | 61.74 ± 5.51 b | 9.20 ± 5.20 a | 29.08 ± 1.20 b | 0.47 ± 0.05 b | 14.24 ± 1.65 b | 0.61 ± 0.02 a |
| August 8, 2017 | Water | 65.27 ± 3.45 b | 14.21 ± 4.42 ab | 20.30 ± 2.29 a | 0.31 ± 0.03 a | 11.93 ± 1.80 ab | 0.63 ± 0.02 a |
| March 5, 2018 | Water | 62.37 ± 3.40 a | 14.60 ± 2.00 a | 22.33 ± 5.25a | 0.36 ± 0.11 a | 11.10 ± 0.62 a | 0.71 ± 0.02 a |
| March 5, 2018 | H<sub>2</sub>CN<sub>2</sub> | 64.50 ± 1.39 ab | 13.83 ± 1.89 a | 21.03 ± 1.76 a | 0.33 ± 0.03 a | 11.40 ± 0.56 a | 0.74 ± 0.01 a |
| May 13, 2018 | Water | 70.50 ± 2.11 b | 9.71 ± 1.84 a | 18.57 ± 1.25 a | 0.26 ± 0.02 a | 12.15 ± 0.64 a | 0.72 ± 0.01 a |
| August 4, 2018 | Water | 72.77 ± 2.50 b | 10.68 ± 2.38 a | 16.57 ± 2.38 a | 0.23 ± 0.03 a | 12.13 ± 1.47 a | 0.79 ± 0.03 a |

Note: G1, S, G2, CV, and debris factor are shown in percent and mitotic index are shown as the ratio of G2 over G1. The mean and standard deviation values are followed by different letters indicate significant differences at $p < 0.05$ according to a Tukey multiple comparison test (n=3).

### Ultrastructure analysis of the grapevine bud shoot apical meristem

The TEM observations were conducted of the shoot primordia of the primary buds collected in March, May, and August 2017. Semi-thin sections at 1 µm were made to trim resin-embedded buds until the area of the shoot primordium was visible (**Figure 3A**). Morphology observations were made at two regions of the SAM, i.e. the central zone (CZ) and peripheral zone (PZ) (**Figure 3B**), and the leaf primordia (LP) (Figure 7A).

**Figure 3.**
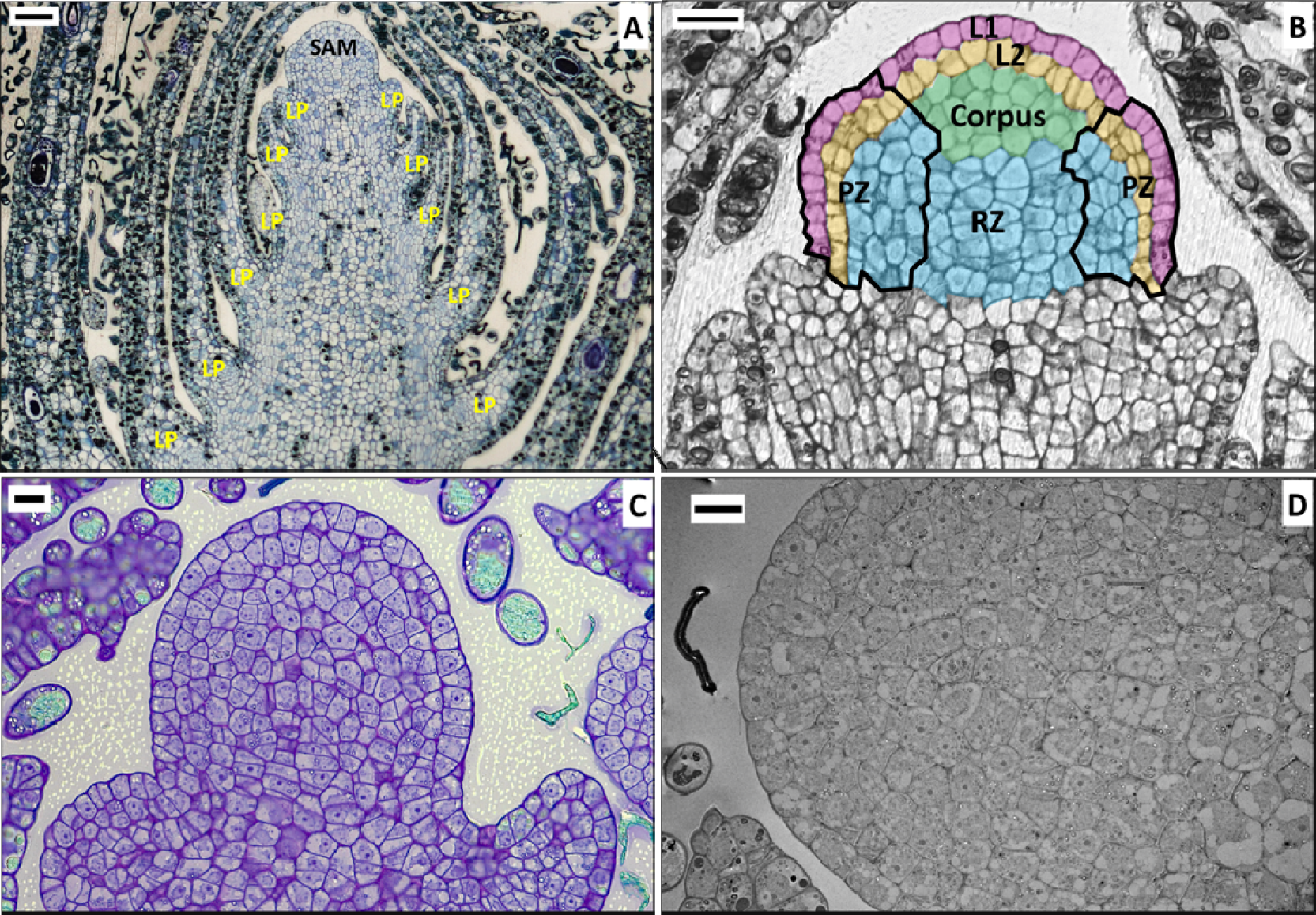
Median longitudinal section of the vegetative shoot of the Vitis vinifera cv. Cabernet Sauvignon primary bud. (A) Light micrograph showing the zone of the shoot apical meristem (SAM) and six leaf primordia (LP)/ nodes. Bar = 50 µm. (B) Schematic interpretation of the SAM zonation according to Mullins et al. (1992) showing the central zone consisting of a tunica (L1 and L2 cell layers) and corpus, peripheral zone (PZ) and rib meristem zone (RZ). Bar = 20 µm. (C) Representative light micrograph of the apical dome of SAM and two leaf primordia. Bar = 10 µm. (D) Representative low magnification electron micrograph of the apical dome. Bar = 10 µm

**Figures 4-7** show representative TEM images of the SAM of grapevine buds at each time or treatment condition in the tunica (**Figure 4**), PZ (**Figure 5**), RZ (**Figure 6**), and LP (**Figure 7**). Starch grains were present in cells throughout the SAM in each time or treatment condition, with relative abundance was higher in March and August than in May buds. The nucleolus and heterochromatin were the most apparent structures in the CZ area. Heterochromatin patches at the periphery of the nuclear envelope were clearly visible in the nucleus of buds collected in March and treated with hydrogen cyanamide (**Figure 4A-B**). A nucleolus can be identified as a densely stained spherical structure inside a nucleus. Meanwhile, the euchromatin usually appeared as a thin fibre and scattered throughout the nucleoplasm, which then began to condense as the cells prepared for cell division, appearing as a small densely stained patch of heterochromatin. The cell walls throughout the SAM of buds collected in March and August appeared thicker than those collected in May. Endoplasmic reticulum was visible in bud cells collected in May (black arrow, **Figure 4E-F**) and relatively absent in bud collected in March (**Figure 4C-D**) and August (**Figure 4G-H**). Numerous osmiophilic organelles were visible (indicated by the asterisk), especially in buds collected in August (**Figure 4-H**); however, no detailed internal membrane structure, e.g. grana or cristae, could be observed, making it difficult to identify these organelles. Cells at the peripheral zone were identified by the presence of tannin deposition (**Figure 5**). The vacuoles were found larger at the rib meristem zone and occupied a significant part of the cells (**Figure 6**) in contrast to cells at tunica, which were largely occupied by the nucleus (**Figure 4**). Ultrastructural details of the nucleus were most resolute in the LP area (**Figure 7A**), showing the double membrane of the nuclear envelope (red arrowheads), nucleolus, and heterochromatin at the periphery of the nucleus (**Figure 7B**). Also visible, the deposition of tannin inside vacuole (**Figure 7B**), starch grain inside a plastid (**Figure 7C**), plastoglobulins inside a proplastid (**Figure 7C**), endoplasmic reticulum (**Figure 7C-D**; black arrowheads), and the cristae membrane in the mitochondria (**Figure 7D**).

**Figure 4.**
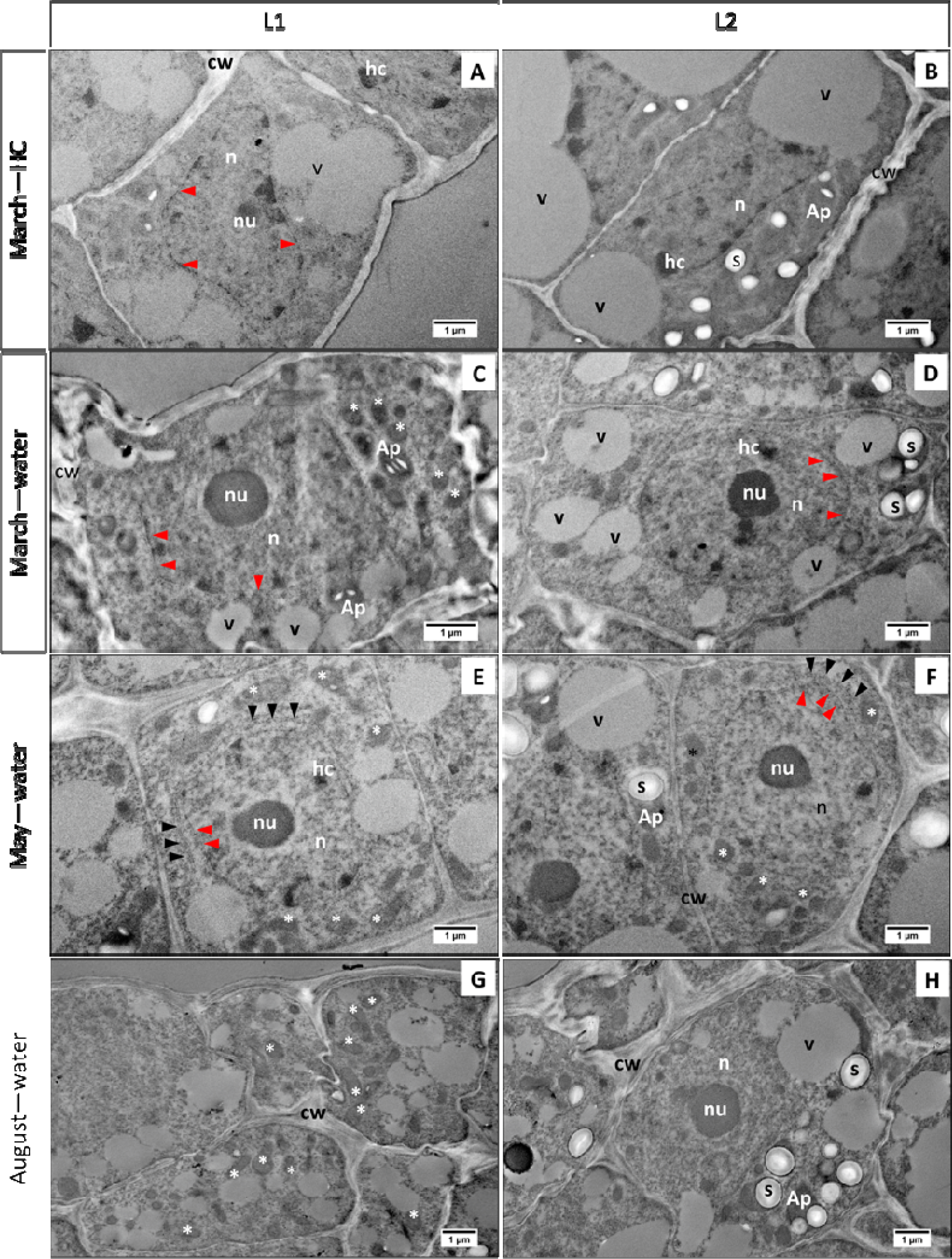
Electron micrograph at the tunica layers (L1 and L2) at the shoot apical meristem. (**A-B**) Tunica of buds collected in March treated with hydrogen cyanamide. Nuclei (n) with nucleoli (nu) and heterochromatin (hc) patches were highlighted at both L1 and L2. Starch grains (s) were visible at L2 and some located inside amyloplast (Ap). (**C-D**) Tunica of buds collected in March treated with water. Large nucleoli (nu) were observed inside the nuclei (n) at L1 and L2. Also visible are few heterochromatin (hc) patches, the double membrane of nucleus envelope (red arrowhead), and numerous small vacuoles (v). Thick cell walls (cw) appeared at L1 of tunica. Asterisk indicates numerous osmiophilic organelle, presumably proplastid or lipid bodies. (**E-F**) Tunica of buds collected in May treated with water. Endoplasmic reticulum (black arrowhead) can be observed. Also, visible nuclei (n), large nucleoli (nu), double membrane of nucleus envelope (red arrowhead), heterochromatin (hc) patches, and small vacuoles (v). Numerous osmiophilic organelles are also visible (asterisk). (**G-H**) Numerous osmiophilic organelles (asterisk) and thick cell walls were highlighted in the tunica layers of buds collected in August treated with water.

**Figure 5.**
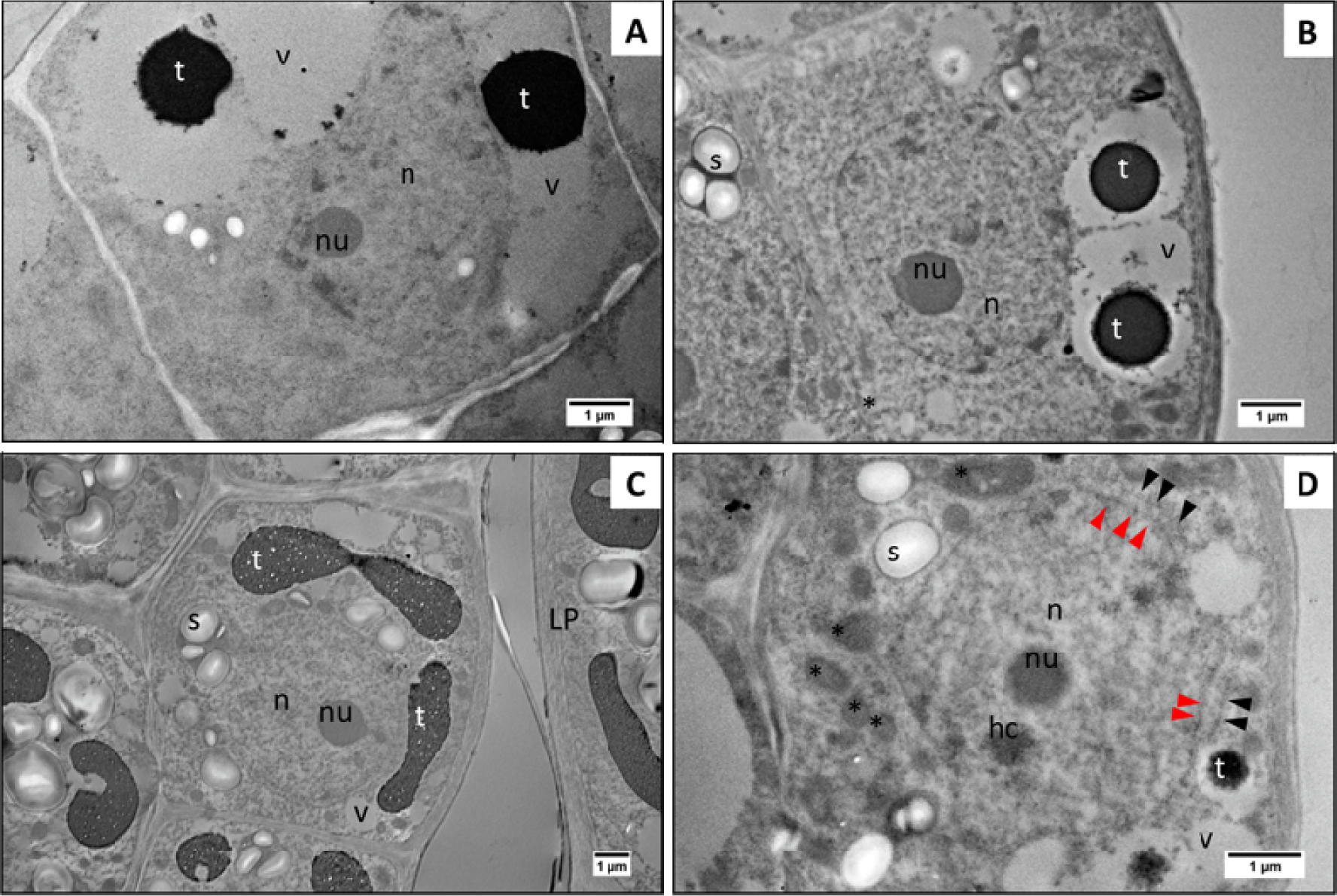
Transmission electron micrograph at the peripheral zone (PZ) of the shoot apical meristem of grapevine buds collected in March (**A**, treated with hydrogen cyanamide; **B**, treated with water), May (**C**), and August (**D**). Tannin depositions were found inside the vacuole of cells at PZ.

**Figure 6.**
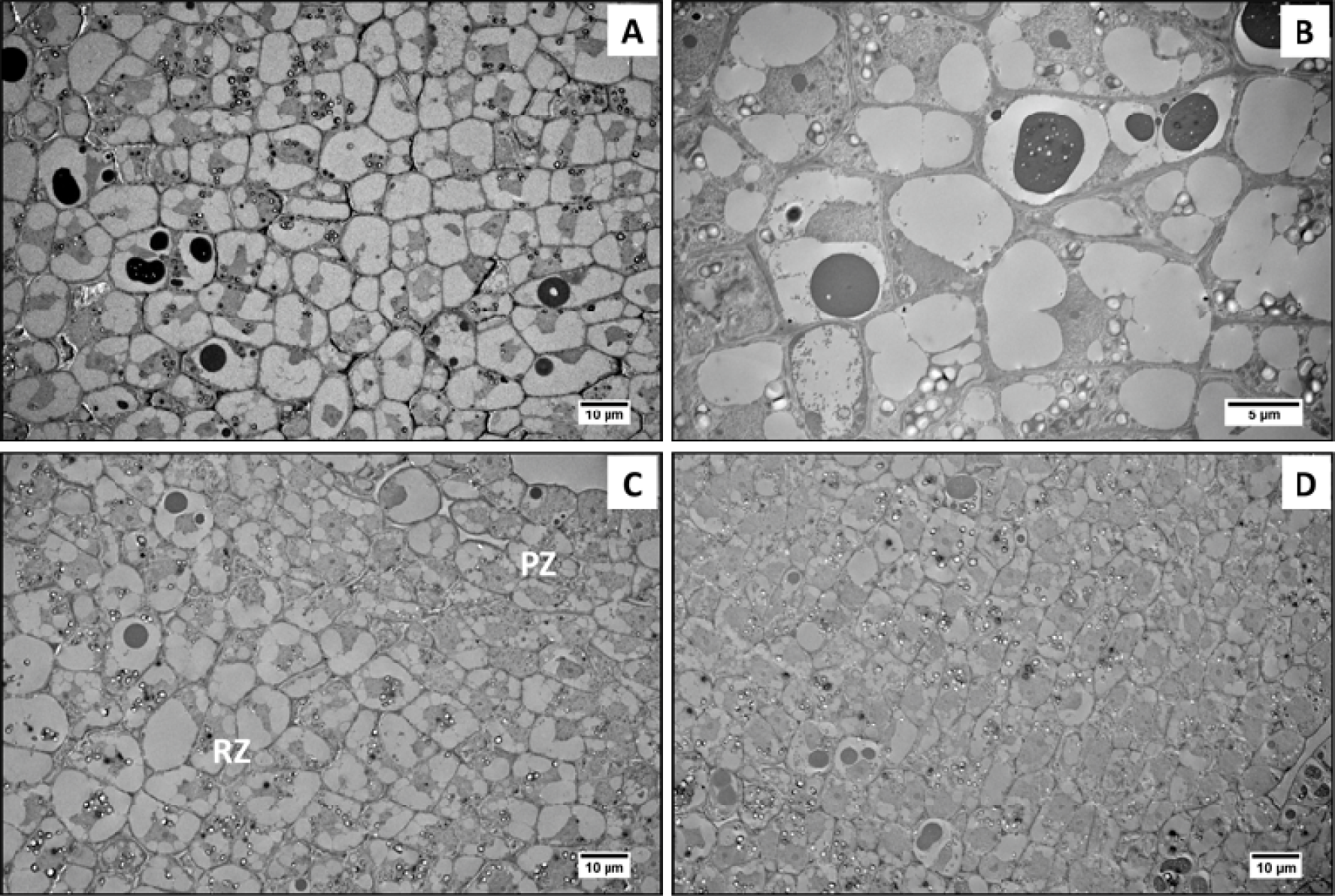
Transmission electron micrograph at the rib meristem zone (RZ) of the shoot apical meristem of grapevine buds collected in March (A, treated with hydrogen cyanamide; B, treated with water), May (**C**), and August (**D**). Cells were relatively larger at RZ compared to PZ mainly due to increase vacuolation. Tannin deposition and starch grains can also be found at this zone.

## DISCUSSION

### Bud cell cycle progression was halted at G1 phase

Several studies have reported changes in nuclear DNA content during seed development and show a different proportion of cells in each cell cycle phase during seed dormancy/ quiescence. This study sought to establish whether we could document parallel changes in nuclear DNA during grapevine bud dormancy. In cabbage seed, only 2C nuclei were detected in dry seed, while 4C DNA was only detected after radicle emerged (Kudo and Kimura, 2010). Time of seed collection and drying process seems to affect the proportion of 2C and 4C DNA in Norway maple seed. The 2C nuclei were dominant in seed harvested after natural shedding (mature and dry), while 4C DNA was found to be more abundant in mature fresh seed (Finch-Savage et al., 1998; Pawłowski et al., 2004). In the axillary buds of Norway maple, dormant buds mainly show 2C nuclear DNA, while 4C DNA was clearly identified in active buds (Bergervoet et al., 1999). Desiccation is suggested to play an important role in the changes of nuclear DNA content during seed development, with a reduction of DNA content coincident with seed maturation and desiccation. Sliwinska (2009) report a higher proportion of 2C nuclei in seed dried at an early stage of maturation and argue that nuclei with a higher DNA content is more sensitive to rapid water loss. Maintaining higher DNA content in the mature dry seed would cause problems later on because of sensitivity to damage, and the presence of damaged DNA will cost time and energy required for DNA repair prior to DNA replication during seed priming and germination. Although indirect, this is an example of a mechanism to protect embryo DNA from degradation under low water content.

In this study, buds collected in March, May and August were mature buds, and it is interesting that the mitotic index does not reflect the depth of dormancy, i.e. the majority of buds have 2C nuclear DNA content or at the G1 phase of cell cycle and the proportion of G1 cells keep increasing towards dormancy release in May and August. If desiccation were suggested to have the same effect on the nuclear DNA of grapevine axillary buds, we would expect that buds collected in May were more desiccated than those collected in March and thus would have more cells with 2C DNA content (G1 phase). As the degree of desiccation in buds (∼40% H_2_O) is not as low as in seeds (∼10% H_2_O), the cell cycle process in dormant buds may not stop completely but slowly progressed and resulted in accumulation of cells arrested in G1 towards August. Despite the convenience of flow cytometry to calculate the proportion of cells in different phases of cell cycles, it does not tell the kinetic of the process. For instance, an increase in the 4C (or endopolyploidy) nuclear content was mostly reported during seed germination process. Compared to bud burst, seeds germination is a relatively quick process and changes in the nuclei content can be observed within 24 hours. This may not be the case with buds. The earliest bud break in non-treated buds was observed on buds collected in August, i.e. 3 weeks after single node planting. It was apparent that cell division in the buds did not resume within 24 hours of treatment, and therefore, the proportion of the nuclei mainly were still at the G1 phase. This was even the case with buds treated with hydrogen cyanamide. An earlier study demonstrated a moderate increase in the expression of *VvCDKA*, *VvCDKB2* and *VvCYCA1*, but not *VvCYCB* or *VvCYCD3.2* in the dormant buds of grapevine within 48 hours of treatment with cyanamide (Vergara et al., 2016). Measuring bud nuclei DNA content at several time points after single node planting (e.g. every 1 week) may show changes in the proportions of nuclei at a different cell cycle phase indicating nuclei DNA content dynamics at the onset of bud burst.

Molecular regulation during dormancy stage is beyond the scope of this study, but several studies reported changes in the expression level of cell cycle related proteins. Both Pawłowski et al. (2004) and Bergervoet et al. (1999) analysed beta-tubulin accumulation analysis on two different embryonic organs, i.e. seed and buds, respectively. Both studies reported that the proportion of G2 never exceeded the proportion of G1. Nevertheless, there was an increase of beta-tubulin protein at the onset of germination/dormancy release. In seeds, beta tubulin accumulation coincides with the decrease of 4C nuclei and increase of 8C nuclei indicating endopolyploidy process instead of mitosis as seeds enter germination. In buds, beta tubulin accumulation increased as the proportion of 4C nuclei in active buds increased. Stanzak et al (2019) report accumulation of ABI5 protein (a transcription factor of the ABA signalling pathway) and RGL2 protein (repressor of GA signalling) at the end of seed development, i.e. mature dry seed. This suggests accumulation is associated with the completion of seed maturation and mainly with desiccation and dormancy acquisition. Investigating the signalling pathway using molecular analysis techniques may provide more depth and insight into the cellular regulation in a slow growing organ such as grapevine buds.

### Subcellular morphology showed the characteristic of cells undergoing dormancy

In the study of annual growth cycle of a perennial tree, knowledge about regulation controlling establishment and release of bud dormancy is less well understood than the regulation of growth cessation and bud set (Singh et al., 2017). Evaluation of dormancy establishment or depth still largely relies on bud response under forcing growth condition (bud burst experiment) which is bearing some limitation, for example standardization of procedure and data analysis (Dennis, 2003; Alvarez et al., 2018). Here, we use mitotic index measurement by flow cytometry and morphology observation using the TEM as an alternative approach to objectively define dormancy stages in grapevine buds. The cellular ultrastructure of SAM region of grapevine buds at all sampling time showed several features in common with the SAM from other plant species at a dormant stage. Cells at the central and peripheral zone of SAM were similar to those observed in *Salix* sp., with the nucleus occupying a significant part of the cells, in addition to undeveloped organelles such as proplastid, and relatively small in size with dense cytoplasm (Berggren, 1985). In contrast, cells at the rib zone are easily identified by the less dense cytoplasm due to increasing vacuolation in the cells and deposition of tannin inside the vacuole (Berggren, 1984). Starch grains were found at the quiescence cells at the central zone of *Heliantus anuus* apical meristem but relatively absent in the more active cells at the peripheral zone (Sawhney et al., 1981). In contrast, starch grains were observed in both the central and peripheral zone of grapevine buds at all time points, indicating that these cells were mitotically less active at the time of sampling. Felker et al. (1983) reported development of starch grains at the SAM of *Prunus cerasus* following chilling application to the quiescent flower buds, whereas quiescent buds without chilling developed more abundant mitochondria and lack of starch grains. We also found similar observation in our experiment, showing fewer starch grains were observed in May buds (quiescent) compared to August buds (quiescent and accumulate chilling). Interestingly in our data, starch grains also appeared relatively abundant in grapevine buds collected in March when the depth of dormancy was the highest and chilling was still absent. It is likely that in grapevine buds, the presence of starch grains indicated two events, i.e. endodormant state and chilling accumulation. Further, the disappearance of starch grain that coincided with the development of mitochondria could only be observed in the more differentiated cells of leaf primordia (**Figure 7**).

**Figure 7.**
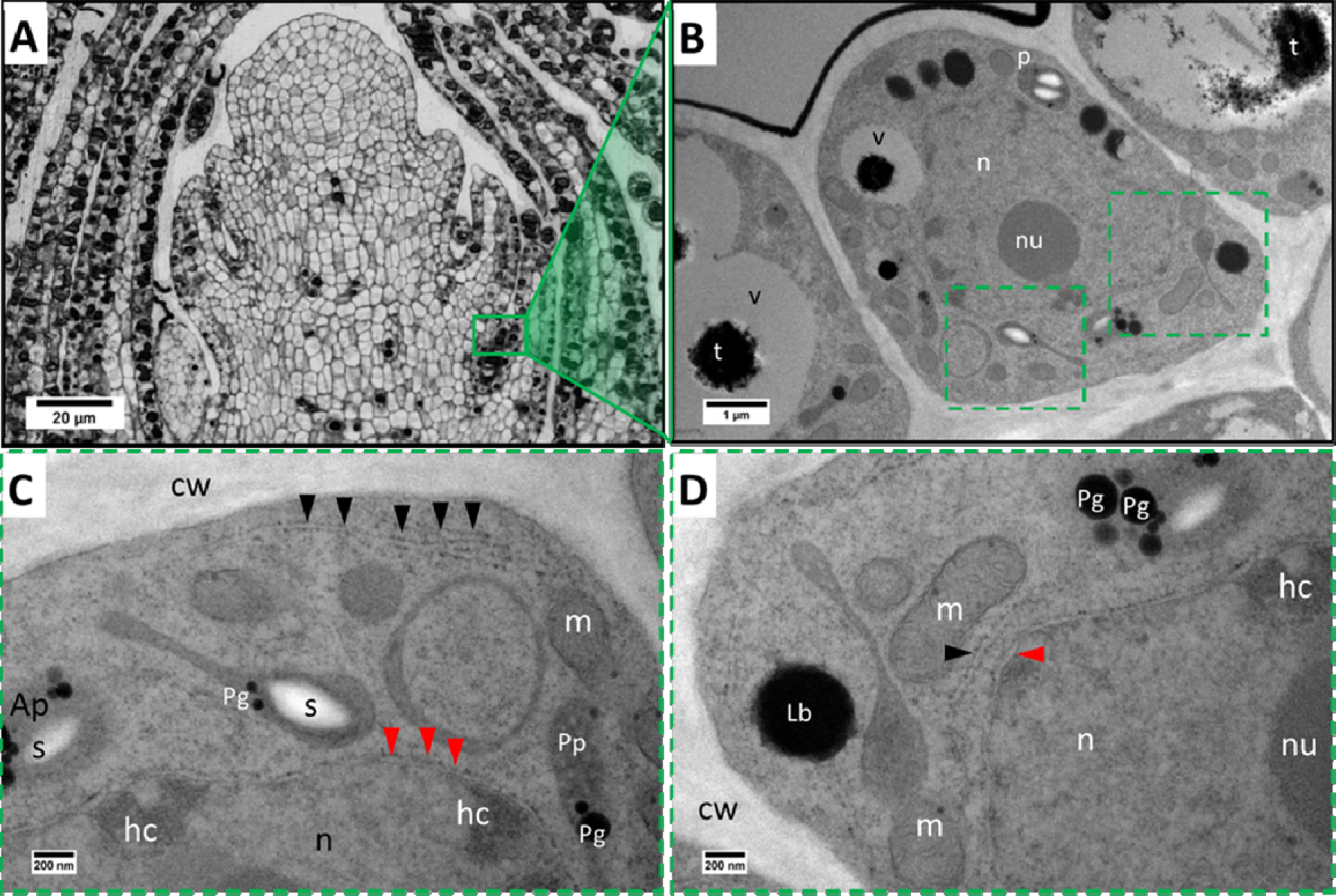
Transmission electron micrograph of cells at the leaf primordia area of mature dormant buds of Vitis vinifera cv. Cabernet Sauvignon revealing cellular ultrastructure of several organelles. (**A**) Light micrograph shoot apex showing the area of observation, i.e., the leaf primordia (red rectangular). (**B**) Representative image of a cell at the leaf primordia area. Visible are nucleus (n), nucleolus (nu), amyloplast (Ap), and vacuole (v) with tannin deposition (t). (**C-D**) Higher magnification images of two areas indicated by dashed rectangular in B. Visible organelles are heterochromatin (hc), proplastid (Pp), amyloplast (Ap), plastoglobulin inside plastid (Pg), osmiophilic organelles presumably lipid bodies (Lb), mitochondria (m), starch grain (s), cell wall (cw). Double membrane nuclear envelope (red arrow heads) and endoplasmic reticulum (black arrow heads) are also visible.

Numerous osmiophilic materials were observed in buds at all time points during the study, which we suggest were either proplastid or lipid bodies. Lipid bodies were commonly found in the shoot apex of both evergreen and temperate perennials (Lynch and Rivera, 1981; van der Schoot et al., 2011; Paul et al., 2014). In the evergreen angiosperm *Rhododendron maximum*, lipid bodies were found to be osmiophilic and mostly restricted to the periphery of the cells (Lynch and Rivera, 1981). The author suggested that lipid bodies act as energy reserves to support metabolic activities. Lipid bodies found in the shoot apex of *Betula pubescens* contain and deliver 1,3-β-glucanase to plasmodesmata region at the plasma membrane and contribute to the removal of plasmodesmatal plug, restoring plasmodesmata function (van der Schoot et al., 2011; Paul et al., 2014). In relation to intercellular communication at the SAM region, plasmodesmata morphology was also found to be regulated during the dormant and active state in *Betula pubescens*. A sphincters structure contains with 1,3-β-D-glucan close the plasmodesmata passage during endodormancy and disappears when the buds are released from dormancy (Rinne et al., 2001). It was suggested that the state of dormancy in the buds was tightly correlated with the regulation of intercellular communications through reversible blockage of plasmodesmata as a response to seasonal changes (Rinne and van der Schoot, 2004). Further, the absence of plasmodesmata sphincters development in transgenic hybrid aspen (*Populus tremula* × *Populus tremuloides* Michx.) was associated with a failure to develop dormancy following exposure to short photoperiod, as bud growth readily resumed in long photoperiod conditions (Tylewicz et al., 2018). The changes of the cell wall thickness at the SAM, as well as presence of starch grain and lipid bodies, were also reported in the SAM of *Cunnighamia lanceolata* during the transition from dormancy to active growth (Xu et al., 2016). In this study, differences in cell wall thickness were more pronounced in the tunica; buds collected in March and August had thicker cell walls than those collected in May. Apart from cell wall thickening, we did not find evidence of plasmodesmata sphincters and lipid bodies in our study.

### Conclusions

With the lack of molecular marker to determine stages of dormancy, i.e., establishment and release, understanding the regulation controlling dormancy in woody perennials is challenging. We coupled bud burst experiment data with mitotic index and ultrastructural morphology of SAM cells data to define the stages of dormancy in grapevine buds. Bud burst experiment showed that peak dormancy of grapevine bud cv. Cabernet Sauvignon was observed during summer when less than 50% buds had burst 200 days after incubation in forcing conditions. Meanwhile, in autumn, the buds attain 50% buds had burst in 60 days, indicating that release from dormancy occurred prior to chilling accumulation. Despite the big difference in the degree of latency on March, May, and August, cell division (G2-M transition) remain absent in the buds at all times of observations and the proportion of cells arrested at G1 keep accumulating toward winter season. The ultrastructure observation of cells at the shoot apical meristem showed an absence of mature organelles at all times of observation and the accumulation of starch in March and August buds associated with dormant and chilling-accumulated quiescent buds, respectively. Our result suggests interesting cellular evidence correspond to the growth resumption capacity of grapevine buds, i.e., absence of mitosis activities regardless of dormancy depth and starch accumulation irrespective of chilling accumulation.

## MATERIALS AND METHODS

### Plant material

Buds of grapevine (*Vitis vinifera* cv. Cabernet Sauvignon) were harvested from c. 100 vines within a commercial cane-pruned vineyard in the temperate/Mediterranean region of Margaret River, Western Australia (34°S, 115°E) on six dates, from early autumn to mid-winter of 2017 and 2018; i.e., 5^th^ March 2017, 16^th^ May 2017, 8^th^ August 2017, 5^th^ March 2018, 13^th^ May 2018, and 4^th^ August 2018. These dates were chosen as preliminary data showed a six-fold range in the depth of dormancy, measured as the time for 50% of buds to burst (BB_50_) to the stage when a green leaf tip become visible or EL4 of the modified Eichorn-Lorenz scale (Coombe, 2004) under forcing conditions (20°C with a 12 hours photoperiod). Canes were cut from the vines within 2 hours from sunrise and trimmed to five nodes from the 4^th^ to the 8^th^ node (1^st^ clear node should be ≥10 mm from the base of the branch). The canes were wrapped in damp newsprint and immediately transported to the lab in an insulated box and stored at 22°C for up to 24 hours. The canes were cut into short single node cuttings of 50-70 mm length, retaining 5-10 mm above the node, as previously described in Meitha et al. (2015). Treatment with hydrogen cyanamide (H_2_CN_2_; Sigma-Aldrich #187364) was done by submerging the node into 1.25% (w/v) H_2_CN_2_ for 30 seconds. Control buds were treated in the same manner with water (W). The cuttings were then stored in the dark for 24 hours before processing and subsequent analysis.

For flow cytometry analysis, tomato (*Lycopersicum esculentum* cv. Money Maker) seeds were germinated and grown in vermiculite. Seedlings were watered daily and maintained at 22°C under cool white LED light and 12 hours photoperiods. The newly emerged young leaves of size no more than 10 mm were harvested from seedlings.

### Depth of dormancy assessment

Fifty single node cuttings per treatment were used to measure the depth of dormancy at each harvest time. The single node cuttings were planted in potting mix (fine composted pine bark, coco peat, and brown river sand in the ratio of 2.5:1:1.5 (w/w) pH∼6.0), in a well-drained seedling tray. Cuttings were grown in a controlled environment room at 20°C, 12 hours photoperiod (450 µmol.m^2^s^1^ Photosynthetically Active Radiation from Eye Hortilux 1000-Watt lamps, Ohio, USA). Trays were watered every 2 days to 80% field capacity. The bud burst (EL4) date was recorded every day for up to 300 days. Non-burst buds were dissected at the end of the assessment period to observe the viability of buds.

### Flow cytometry (FCM)

Nuclei preparation, FCM data acquisition, and data analysis were performed as described by Hermawaty et al. (2022). FCM using buds was performed in three biological replicates in which ten buds were pooled in each biological replicate. Two canes from two independent vines with 5 buds each were used to make a pool of ten primary buds for one replicate. The single bud is the biological unit of interest but considering that nuclei represent a minority population within the nuclei suspension, therefore in this experiment, it was necessary to pool the buds to obtain a sufficient number of nuclei for FCM analysis.

Primary buds were manually dissected under a dissecting microscope, carefully removing the secondary and tertiary buds and woolly-hairs as much as possible. A total of ten primary buds (± 100 mg) per replicate were collected into a plastic petri dish (diameter, 100 mm) and placed on ice. Nuclei was extracted by individually chopped sample using a sharp razor blade in 3 mL cold nuclear isolation buffer (NIB; 10 mM MgSO4.7H2O, 50 mM KCl, 5 mM HEPES, and 1 mg.mL^-1^ DTT, 0.5% Triton X-100, and 3% PVP-40 (w/v)) followed by an hour incubation on ice. After incubation, samples were filtered through a 100 µm and 41 µm nylon mesh filter (Millipore, Germany), respectively, centrifuged at 450×g for 8 minutes at 4°C. Nuclei pellet was resuspended in fresh 1 mL NIB followed by the addition of 50 µg.mL^-1^ of RNase A (Thermo Scientific, Australia, EN0531) to remove RNA from nuclei suspension. After RNase treatment, a final concentration of 20 µg.mL-1 propidium iodide (PI; Sigma-Aldrich #P4170) was added to stain the nuclei. The nuclei suspension without the addition of PI served as the negative control. A mixed nuclei suspension sample was prepared for instrument setting by co-chopping 50 mg of buds (prepared by dissection as described above) with 25 mg of tomato leaves.

The nuclear DNA content was measured using a BD FACSCanto™ II (BD Biosciences, San Jose, CA) flow cytometer equipped with an air-cooled 488-nm solid state 20 mW laser. Propidium iodide was excited using blue laser light (488 nm) and the fluorescent signal was collected through a 556-dichroic long-pass filter and a 585/42 nm band-pass filter. The analysis was run at low speed mode to collect up to 15,000 single PI positive nuclei per biological replicate. The percentage of cells at each phase was estimated using the Watson pragmatic algorithm in FlowJo V10 (Tree Star Inc., Ashland, OR) software. The quality of the nuclei preparation was evaluated using a coefficient of variance (CV), background debris factor (DF), and percentage of intact nuclei as previously described (Hermawaty et al., 2022).

### Ultrastructure observation of the primary buds

Using transmission electron microscopy (TEM), sample preparation and observation were performed at the Centre for Microscopy, Characterisation, and Analysis (CMCA), University of Western Australia. Following the treatment with H_2_CN_2_ or water, as described in the ‘plant material’ section, buds were excised from the canes then immersed in 1.6% formaldehyde and 2.5% glutaraldehyde in phosphate buffered saline (PBS) pH 7.4 solution. Infiltration of the fixative solution was done by applying 2500 mmHg vacuum cycle for 15 minutes (5 min vacuum/release/mixing) at room temperature using a PELCO® vacuum system fitted in the PELCO Biowave® microwave followed by incubation at room temperature for 24 hours (without vacuum). Fixed buds were kept at 4°C until required. Prior to post-fixation with osmium tetroxide, fixed buds were further dissected for removal of secondary and tertiary buds, bud scales and trichomes (**Supplementary Figure S1**). Post-fixation, dehydration, resin infiltration and sample embedding were performed as described by Clode (2015; **Supplementary File 1**). The microscopic study was done on median longitudinal sections (transverse relative to the cane, **Supplementary Figure S1**) of the zone of the shoot apical meristem of the buds. Semi-thin sections at 1 µm were prepared using the resin-embedded buds using a glass knife on an EM UC6 ultramicrotome (Leica Microsystems GmbH, Wetzlar, Germany). Sections were mounted on glass slides, stained with toluidine blue (pH 9; 0.5% toluidine blue in 1% borax) and observed under a bright-field microscope for general morphology. Bright-field images were acquired using an Axioskop optical microscope (Zeiss) fitted with an Axiocam digital camera (Zeiss). Ultrathin sections at 100 nm were prepared using a 45° Histo diamond knife (Diatome, PA, USA) on an EM UC6 ultramicrotome (Leica Microsystems GmbH). Sections were mounted into unfilmed 75-mesh copper grids (ProSciTech, Australia) and imaged unstained at 120 kV using a JEM-2100 TEM (JEOL, Tokyo, Japan). Images were acquired on an ORIUS SC1000 digital camera (Gatan, Pleasanton, CA, USA).

## Abbreviations

SAM: shoot apical meristem
CZ: central zone
PZ: peripheral zone
RZ: rib zone
LP: leaf primordia
BB50: 50% of bud to burst
EL4: Eichorn-Lorenz scale 4
H2CN2: hydrogen cyanamide
FCM: flow cytometry
NIB: nuclei isolation buffer
PI: propidium iodide
CV: coefficient of variance
DF: debris factor
TEM: transmission electron microscopy

## Supplementary data

Supplementary File S1. Sample preparation and processing for general morphology and ultrastructure of grapevine bud using light and transmission electron microscopy.

Supplementary Figure S1. View of mature compound grapevine buds. The compound consists with one primary latent bud which will produce the next season shoot (N+2), and two secondary latent buds (N+3) which only grow when the primary bud die. Two cutting planes relative to the canes are shown above. (1) Longitudinal section in which cutting were made crossing all three buds (2) Transversal section (relative to cane) in which cutting made parallel to the two secondary buds. The median longitudinal view of the shoot apical meristem as shown in Figure 7A is achieved by making transversal cutting relative to the cane.

## Acknowledgements

We are thankful to Keith Mugford of Moss Wood Wines for enduring support and access to plant material at often inconvenient times. We acknowledge the facilities, and scientific and technical assistance of Microscopy Australia at the Centre for Microscopy, Characterisation & Analysis, The University of Western Australia. We also acknowledge support and teamwork of other laboratory members, particularly Juwita Dewi.

## Author contributions

DH: planned, performed all the experiments, analysed the data, prepared all the figures, and wrote the manuscript with constructive comments from MJC. PLC, SS and JAC: assisting with experimental design, data acquisition, and interpretation of microscopy analysis. MJC: conceived and supervised the project. All authors contributed to the article and approved the submitted version.

## Conflicts of interest

The authors declare no conflict of interests.

## Funding

The authors acknowledge funding support of the Australian Research Council (DP150103211, FT180100409). Microscopy Australia at the Centre for Microscopy, Characterisation & Analysis, is funded by the University, State and Commonwealth Governments.

## Data availability

Data are available from the corresponding author, Michael Considine, upon request.

## Notes

### Competing Interest Statement

The authors have declared no competing interest.

